# Unveiling fetal brain changes in congenital diaphragmatic hernia: hypoxic injury with loss of progenitor cells, neurons, and oligodendrocytes

**DOI:** 10.1101/2023.09.23.559137

**Authors:** George Biouss, Lina Antounians, Julien Aguet, Katarina Kopcalic, Nikan Fakhari, Jerome Baranger, Luc Mertens, Olivier Villemain, Augusto Zani

## Abstract

Congenital diaphragmatic hernia (CDH) is a birth defect characterized by incomplete closure of the diaphragm, herniation of abdominal organs into the chest, and compression of the lungs and the heart. Besides complications related to pulmonary hypoplasia, 1 in 4 survivors develop neurodevelopmental impairment, whose etiology remains unclear. Using a fetal rat model of CDH, we demonstrated that the compression exerted by herniated organs on the mediastinal structures results in decreased brain perfusion on ultrafast ultrasound, cerebral hypoxia with compensatory angiogenesis, mature neuron and oligodendrocyte loss, and activated microglia. In CDH fetuses, apoptosis was prominent in the subventricular and subgranular zones, areas that are key for neurogenesis. We validated these findings in the autopsy samples of four human fetuses with CDH compared to age- and sex-matched controls. This study reveals the molecular mechanisms and cellular changes that occur in the brain of fetuses with CDH and creates opportunities for therapeutic targets.

## Introduction

Congenital diaphragmatic hernia (CDH) is a severe birth defect characterized by the incomplete formation of the diaphragm which causes herniation of fetal intra-abdominal organs into the chest (1). The compression exerted by the herniated organs onto the intra-thoracic structures contributes to pathological changes observed in the lung (pulmonary hypoplasia with impaired branching morphogenesis and aberrant vascularization) and heart (left ventricular hypoplasia) (1–3). These cardiopulmonary developmental anomalies are responsible for the high mortality (25-30%) and morbidity (>60%) rates that still burden babies with CDH (1, 4). Moreover, 1 in 4 CDH survivors suffers from neurodevelopmental impairment (NDI) that may affect any one of the domains, ranging from gross and/or fine motor and sensory (hearing, visual) deficits to cognitive, language, and behavioral impairment. One in 2 survivors have abnormal findings at postnatal neuroimaging (5).

The etiology of NDI in CDH is unknown. In patients with isolated CDH, i.e., those with no genetic anomalies or severe co-morbidities, NDI is generally ascribed to invasive postnatal interventions, such as prolonged mechanical ventilation and/or extracorporeal life support (1). However, there is growing evidence that an insult to the brain may occur before birth and predispose to postnatal NDI. A retrospective review of fetal ultrasound exams showed that fetuses with prenatally diagnosed CDH had lower middle cerebral artery (MCA) flow velocity compared to normal controls (6). Another two studies using fetal ultrasound scans confirmed alterations in MCA velocity, which was found to be severely decreased in the third trimester and to correlate with a decreased left ventricular cardiac output on fetal echocardiogram (7, 8). Moreover, antenatal ultrasound and magnetic resonance studies showed that the alterations in brain perfusion observed in fetuses with CDH were associated with lower fetal brain volumes, especially in the cortex, hippocampus, corpus callosum and cerebellum (7, 9–11). However, due to the limitations of the imaging techniques and the unavailability of brain tissue data, these studies could not determine the changes that occur in the brain tissue of fetuses with CDH.

To further investigate the molecular mechanisms and cellular changes involved in brain injury in CDH fetuses, we conducted a study based on a fetal rat model and validated our findings using autopsy samples from human fetuses with CDH. With our model, we showed that the brain of rat fetuses with CDH has decreased global and regional cerebral blood perfusion and shows signs of hypoxia with compensatory angiogenesis. Moreover, we found increased apoptosis of the progenitor cell populations in the brain stem cell niches, associated with decreased density of neurons and oligodendrocytes proportional to the degree of brain hypoxia. Brain autopsy specimens from human fetuses with CDH had similar features of cerebral hypoxic injury with loss of progenitor cells, neurons, and oligodendrocytes. Taken together, these findings increase our understanding of the pathophysiology of NDI predisposition in CDH and create opportunities for therapeutic targets.

## Results

### The brain of rat fetuses with CDH has altered perfusion with evidence of hypoxia and compensatory angiogenesis

To study the molecular mechanisms and cellular changes in the brain of CDH fetuses, we employed the most widely used model based on nitrofen administration to rat dams at embryonic day (E)9.5 (12, 13). It is well established that maternal administration of nitrofen at this time point results in pulmonary hypoplasia alone (no CDH) in about half of the litter and in pulmonary hypoplasia with CDH (diaphragmatic defect with intra-abdominal organ herniation generating mechanical compression of the intra-thoracic organs) in the rest of the litter (12). As we hypothesized that cerebral flow impairment in CDH is due to the compression of mediastinal structures, we used this model where nitrofen-exposed rat fetuses without CDH served as an internal control.

We first elected to establish whether this rat model of CDH recapitulated the decreased brain perfusion observed in human fetuses with CDH (6, 7). To this end, we scanned the brain of rat fetuses using ultrafast ultrasound power doppler imaging *in vivo*. This is a novel ultrasound technique based on plane-wave imaging that quantifies cerebral blood volume (CBV) (14, 15). Compared to healthy controls, the CBV of the whole brain (global CBV) was decreased by 29% in nitrofen-exposed fetuses without CDH and by 61% in fetuses with CDH (Figure 1, A and B). Moreover, nitrofen-exposed fetuses with CDH had lower global CBV than nitrofen-exposed fetuses without CDH (71% vs. 39% relative to control, p=0.002). When we focused on the middle cerebral region that is known to be affected in human fetuses with CDH (8), we observed that fetuses with CDH had a 55% reduction in CBV compared to healthy controls (p=0.002) and 32% compared to nitrofen-exposed fetuses without CDH (p=0.048; Figure 1, A and B). In the anterior cerebral region, there were no differences in CBV between nitrofen-exposed fetuses with and without CDH (Supplementary Figure 1), whereas using ultrafast ultrasound imaging, it was not technically feasible to detect flow in the posterior cerebral region.

**Figure 1.**
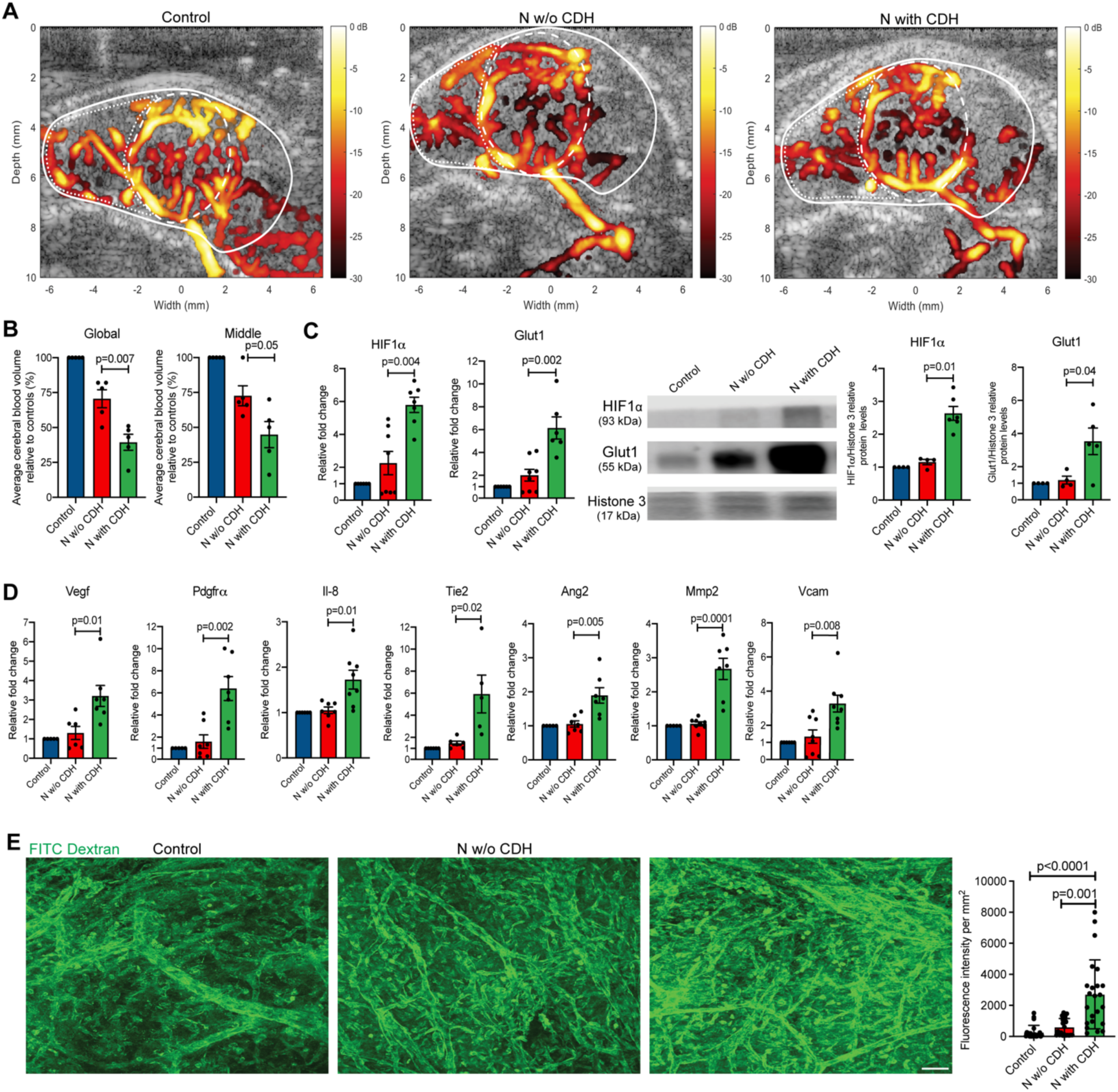
Experimental CDH is associated with lower cerebral blood volumes resulting in hypoxia and compensatory angiogenesis. **A** Representative image obtained from ultrafast ultrasound doppler scans of rat fetal brains at embryonic day E21.5 in three conditions. Solid white line indicates global cerebral area, dashed line indicates middle cerebral area. **B** Average cerebral volume quantification in the global and middle cerebral relative to control fetuses from n=5 biological replicates in each condition. **C** Gene and protein expression of hypoxia response genes including *HIF1α* and *Glut1* for Control (n=6), N w/o CDH (n=8), and N with CDH (n=6) for RNA expression and Control (n=4), N w/o CDH (n=5), and N with CDH (n=5) for protein expression. **D-G** Gene expression of vascular remodeling and angiogenesis markers in three conditions for Control (n=5), N w/o CDH (n=6), and N with CDH (n=6). **H** Representative images obtained from injection of FITC, quantified by fluorescence intensity as a proxy for cerebral vascular density from at least n=22 fields per condition. Scale bar = 100 μm. Data are presented as mean ± SD.

To determine whether the changes in CBV observed in rats with CDH had an impact on brain oxygenation, we measured hypoxia markers in the fetal rat brain. We observed that nitrofen-exposed rat fetuses with CDH had higher gene and protein expression levels of canonical markers of hypoxia (HIF1α, Glut1) compared to both healthy controls and nitrofen-exposed fetuses without CDH (Figure 1C). Importantly, we did not detect any differences in hypoxia markers between healthy controls and nitrofen-exposed fetuses without CDH (Figure 1C). We then investigated whether the brain of fetuses with CDH undergoes compensatory angiogenesis, an adaptive mechanism aimed at optimizing oxygen delivery to tissues. Compared to both healthy controls and nitrofen-exposed fetuses without CDH, the brain of nitrofen-exposed fetuses with CDH had higher gene expression levels of pro-angiogenic growth factors and chemokines (*Vegf, Pdgfra, Il-8*), endothelial cell proliferation factors (*Tie1, Ang2*), and markers of extracellular matrix turnover (*Mmp2, Vcam*) (Figure 1, D and E). No differences were found in all these factors between healthy controls and nitrofen-exposed fetuses without CDH (Figure 1D). To corroborate these findings, we also measured the density of cerebral vessels by labeling them with fluorescein isothiocyanate (FITC)-dextran and imaging them with two-photon laser microscopy. We observed that the brain of nitrofen-exposed CDH fetuses had higher vascular density compared to healthy controls and nitrofen-exposed fetuses without CDH (Figure 1E). The latter two groups had no differences in vascular density. Overall, these data confirmed that the presence of CDH is associated with brain hypoxia and compensatory vascular adaptation.

### Neuron and oligodendrocyte progenitor cell populations in the subventricular and subgranular zones have increased levels of apoptosis in the brain of rat fetuses with CDH

Given the perfusion changes observed in the brain of rat fetuses with CDH, we first examined the gross morphometry of the brain, and then investigated the biological processes that are typically affected by tissue hypoxia, such as apoptosis, endoplasmic reticulum stress, and lipid peroxidation (16–18). On gross morphometry, we found that nitrofen-exposure caused a reduction in brain size, weight, and brain/body weight ratio regardless of the presence of CDH (Figure 2A). When we assessed apoptosis, we found that the brain of nitrofen-exposed fetuses with CDH had higher gene and protein levels of pro-apoptosis markers compared to nitrofen-exposed fetuses without CDH and healthy controls (Figure 2A, Supplementary Figure 2). To identify which regions of the brain had higher levels of apoptosis, we performed terminal deoxynucleotidyl transferase–mediated deoxyuridine triphosphate nick end labeling (TUNEL) staining on whole brain sections. We found the highest number of apoptotic cells in the subventricular zone (SVZ) and subgranular zone (SGZ), which are neural stem cell niches (Figure 2B). In these regions, apoptosis of neural precursor cells (NPCs) and oligodendrocyte progenitor cells (OPCs) was increased in CDH fetuses compared to nitrofen-exposed fetuses without CDH and healthy controls (Figure 2, C and D). We also found that compared to nitrofen-exposed fetuses without CDH and healthy controls, the brain of nitrofen-exposed fetuses with CDH had higher levels of Bip, Eif2s1, and Atf6, indicating an increased endoplasmic reticulum stress response, as well as higher levels of malondialdehyde, an indirect measure of oxidative stress (Figure 2E).

**Figure 2.**
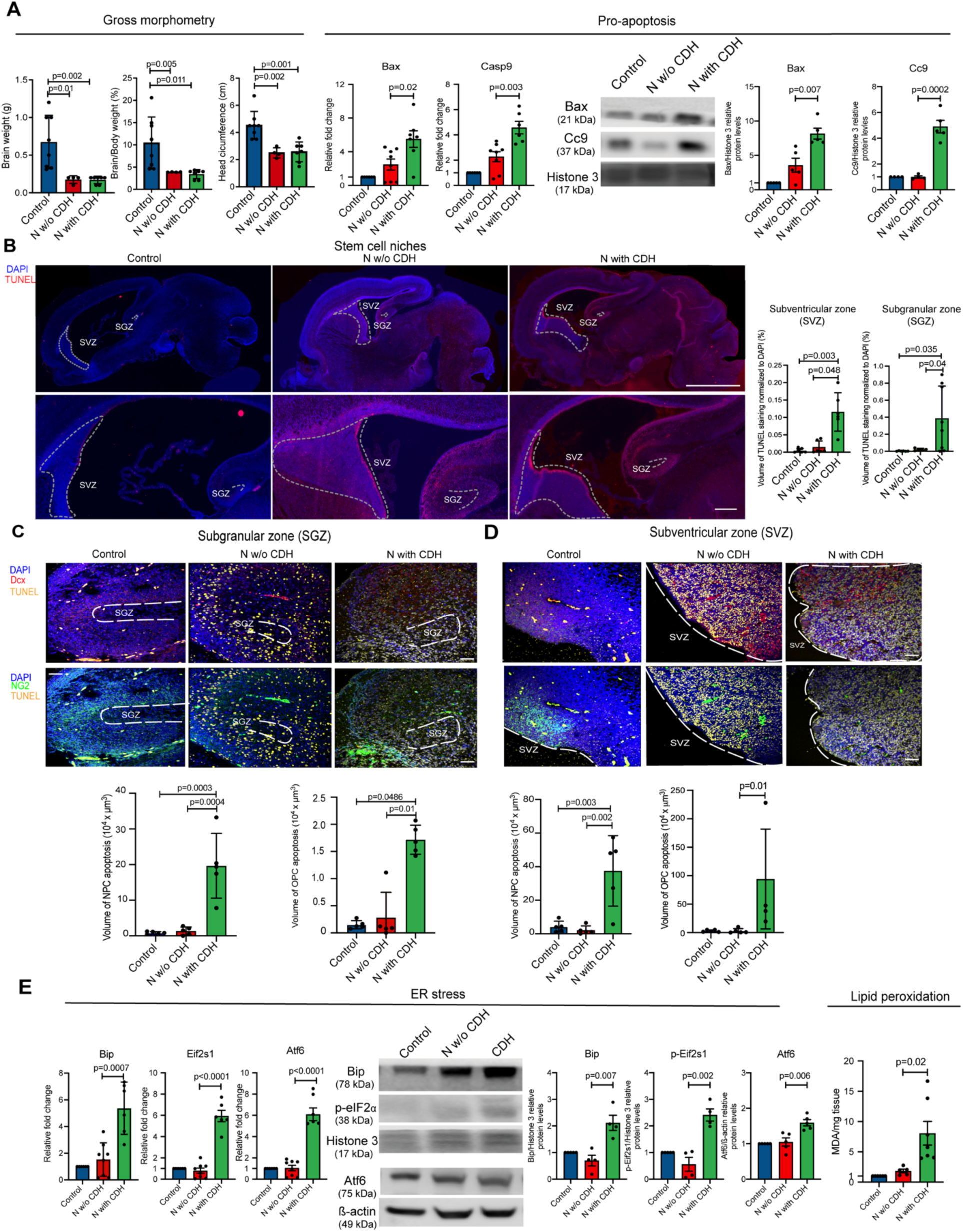
Neuron and oligodendrocyte progenitor cell populations in the subventricular and subgranular zones have increased levels of apoptosis in the brain of rat fetuses with CDH. **A** Gross morphometric measurements of rat fetal brain weights, brain-to-body weight ratio, and head circumference from Control (n=8), N w/o CDH (n=4), and N with CDH (n=7). Gene and protein expression of pro-apoptotic factors BCL2 Associated X (Bax) and Caspase-9 (Casp9). **B** Representative immunofluorescence images of terminal deoxynucleotidyl transferase–mediated deoxyuridine triphosphate nick end labeling (TUNEL) assay (red) and nuclear marker (DAPI, blue) in sagittal sections of rat fetal brains. Dotted lines indicate borders of subgranular zone (SGZ) and subventricular zone (SVZ) from Control (n=5), N w/o CDH (n=5), and N with CDH (n=5). Quantification of total TUNEL volume per nuclear region derived from multiple fields merged into a Z-stack is shown. **C-D** Representative co-staining immunofluorescence images of markers of apoptosis (TUNEL, yellow), neuronal precursor cells (NPC) doublecortin (DCX, red), oligodendrocyte progenitors neural/glial antigen 2 (NG2, green), and nuclear stain (DAPI, blue), in the SGZ and SVG. Quantification of NPC and OPC apoptosis is shown. Data are presented as mean ± SD. **E** Gene and protein expression of endoplasmic reticulum stress response factors. Control (n=6), N w/o CDH (n=8), and N with CDH (n=6). Reactive oxygen species activity measured by the degree of lipid peroxidation (TBAR assay). The ratio of lipid peroxidation product malondialdehyde/total protein yield (mg) was measured. Control (n=6), N w/o CDH (n=5), and N with CDH (n=7).

### Cerebral hypoxia in rat fetuses with CDH is associated with reduced mature neuron, oligodendrocyte density, and microglia activation

The high levels of apoptosis affecting NPCs and OPCs led us to interrogate whether the number of mature neurons and oligodendrocytes was also reduced. To this end, we focused on the areas of the brain that were reported to be affected in human prenatal imaging studies, that is the cerebral cortex, hippocampus, and cerebellum (7, 9–11). In the cerebral cortex, we found that fetuses with CDH had significant reduction in mature neuron density and proliferating oligodendrocytes (Figure 3A). Moreover, nitrofen-exposed fetuses with CDH had fewer mature neurons and proliferating oligodendrocytes compared to nitrofen-exposed fetuses without CDH (Figure 3A). In the hippocampus, fetuses with CDH also had lower density of mature neurons and lower proliferating oligodendrocytes compared to healthy controls and with nitrofen-exposed pups without CDH (Figure 3B). Lastly, in the cerebellum, nitrofen-exposed fetuses with CDH also had the lowest number of mature neurons and proliferating oligodendrocytes compared with both healthy controls and nitrofen exposed pups without CDH (Figure 3C). We also assessed the density of microglia and astrocytes in the same brain regions. Activated microglia had the highest density in the hippocampus and cerebellum of nitrofen-exposed fetuses with CDH compared to healthy controls and nitrofen-exposed fetuses without CDH (Figure 3, A-C). Astrocyte density was increased only in fetuses with CDH in the hippocampus compared to healthy controls (Supplementary Figure 3).

**Figure 3.**
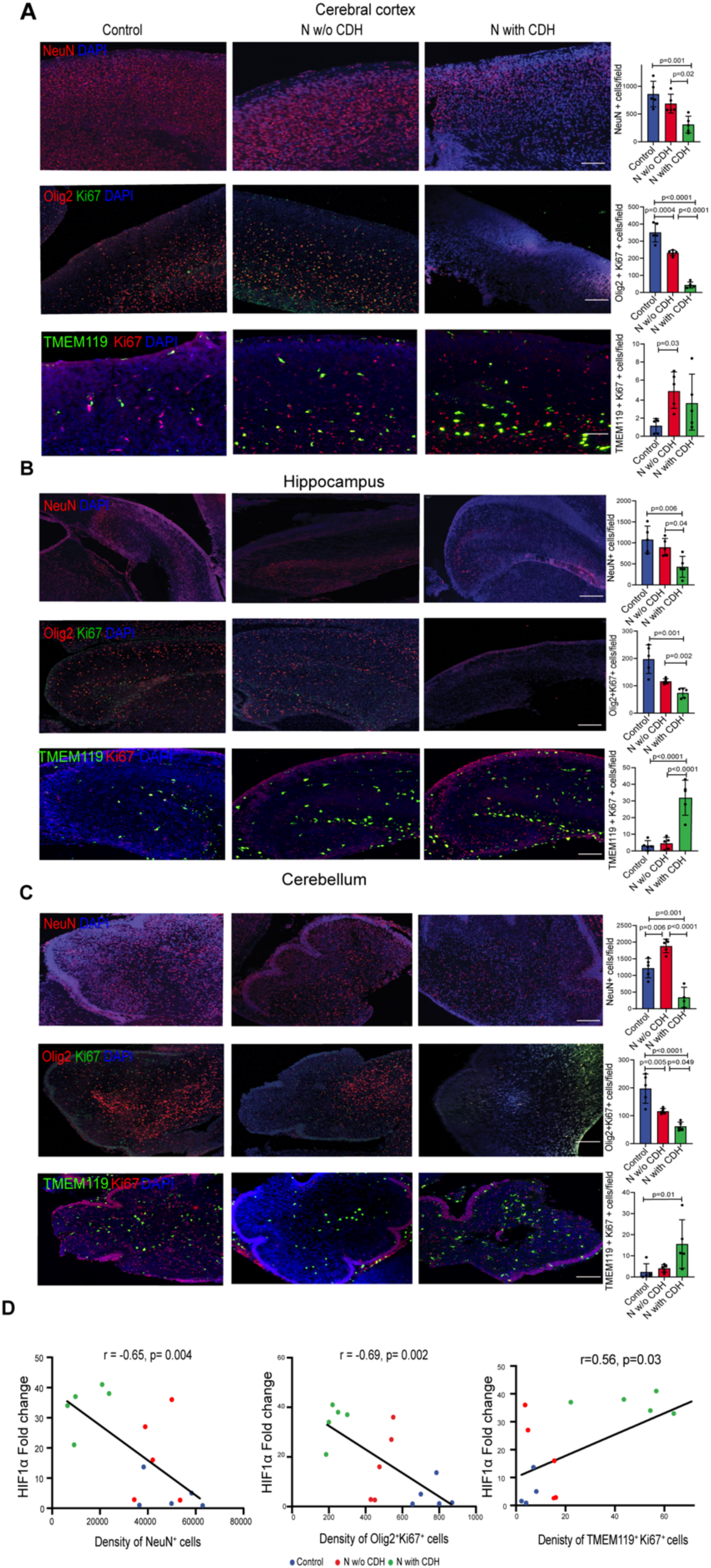
Cerebral hypoxia in rat fetuses with CDH is associated with reduced mature neuron and oligodendrocyte density and microglia activation. **A-C** Representative immunofluorescence images depicting mature neurons (NeuN+), proliferating oligodendrocytes (Olig2+ Ki67+), and activated microglia (TMEM119+ Ki67+) in the cerebral cortex, hippocampus, and cerebellum of rat fetuses. Quantification is conducted as number of cells per field. Scale bar = 50 μm. Control (n=5), N w/o CDH (n=5), N with CDH (n=5) **D** Correlation analysis between *HIF1*α gene expression and mature neurons (left), proliferating oligodendrocytes (middle), or proliferating microglia (right). Pearson correlation statistics are shown. Data are presented as mean ± SD.

We found a negative correlation between the expression of *Hif1α*, a master regulator of hypoxia, and neuronal and oligodendrocyte count, as well as a positive correlation with the number of activated microglia in the same samples (Figure 3D). Specifically, fetuses with CDH had the highest *Hif1α* expression, the lowest density of mature neurons and oligodendrocytes, and the highest number of activated microglia (Figure 3D). These findings suggest that brain cell density changes were influenced by the degree of hypoxia.

### Human fetuses with CDH have cerebral hypoxia with NPC and OPC apoptosis and decrease in mature neurons and oligodendrocytes

To validate our findings of cerebral hypoxia and altered density of cell populations in the fetal rat model of CDH, we examined autopsy samples from the lungs and available brain regions (midbrain and cerebellum) of four fetuses with left-sided CDH, who died or were terminated between 19 and 27 weeks of gestation. Four age- and sex-matched fetuses with no lung or brain pathology served as controls. Demographics and clinical details of the eight fetuses are shown in Table 1. To validate the observations made in the fetal rat model of CDH, we investigated the degree of brain tissue hypoxia, NPC and OPC apoptosis, and the density of mature brain cell populations in the human brain autopsy samples. Compared to controls, human fetuses with CDH had higher levels of activated HIF1α in both the midbrain and cerebellum (Figure 4, A and B). Apoptosis levels of NPCs were higher in both SGZ and SVZ in CDH fetuses compared to controls (Figure 4, C and D). Conversely, we found no differences in the number of apoptotic OPCs in SGZ and SVZ between CDH and control fetuses (Figure 4, C and E). Furthermore, fetuses with CDH had fewer differentiated neurons in the cerebellum and fewer myelinating oligodendrocytes in both midbrain and cerebellum (Figure 4, F and G). No differences were observed in the number of microglia or astrocytes in either region (Supplementary Figure 4).

**Figure 4.**
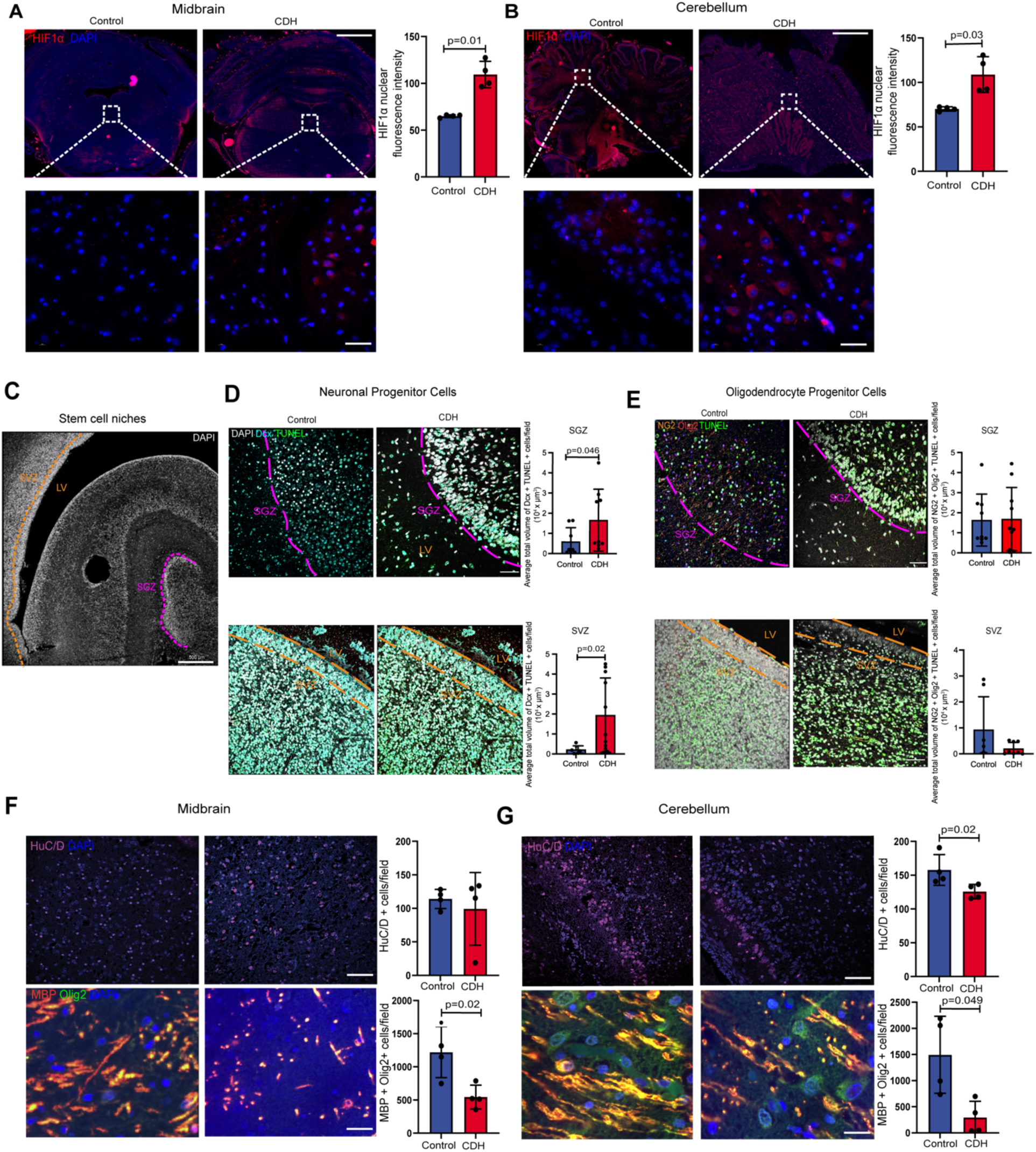
Human fetuses with CDH have cerebral hypoxia with apoptosis of neuron and oligodendrocyte progenitor cell populations and decrease in mature neurons and oligodendrocytes. **A-B** Representative immunofluorescence images in the brain of 4 CDH fetuses and 4 age- and sex-matched controls, in the midbrain (**A**) and cerebellum (**B**) for HIF1α nuclear localization (red; DAPI, blue), HuC/D oligodendrocytes (purple; DAPI, blue), and myelin basic protein (MBP, red) and mature oligodendrocytes (Olig2, green; DAPI, blue). Quantification is shown as fluorescence intensity (HIF1α). Scale bar = 50 μm. **C** Representative immunofluorescence image of subventricular zone (SVZ), subgranular zone (SGZ), and lateral ventricles (LV) in human fetal autopsy samples (DAPI, white). **D** Representative immunofluorescence images of apoptotic cells (TUNEL+, green), NPCs (DCX, blue), and nuclear marker (white), and **E** OPCs (NG2/Olig2) in fetal CDH autopsy samples and controls. Quantification shown as total volume of cells per field (multiple Z-stacks). **F** Number of cells per field in the midbrain (HuC/D, MBP/Olig2) **G** Number of cells per field in the cerebellum.

**Table 1.**
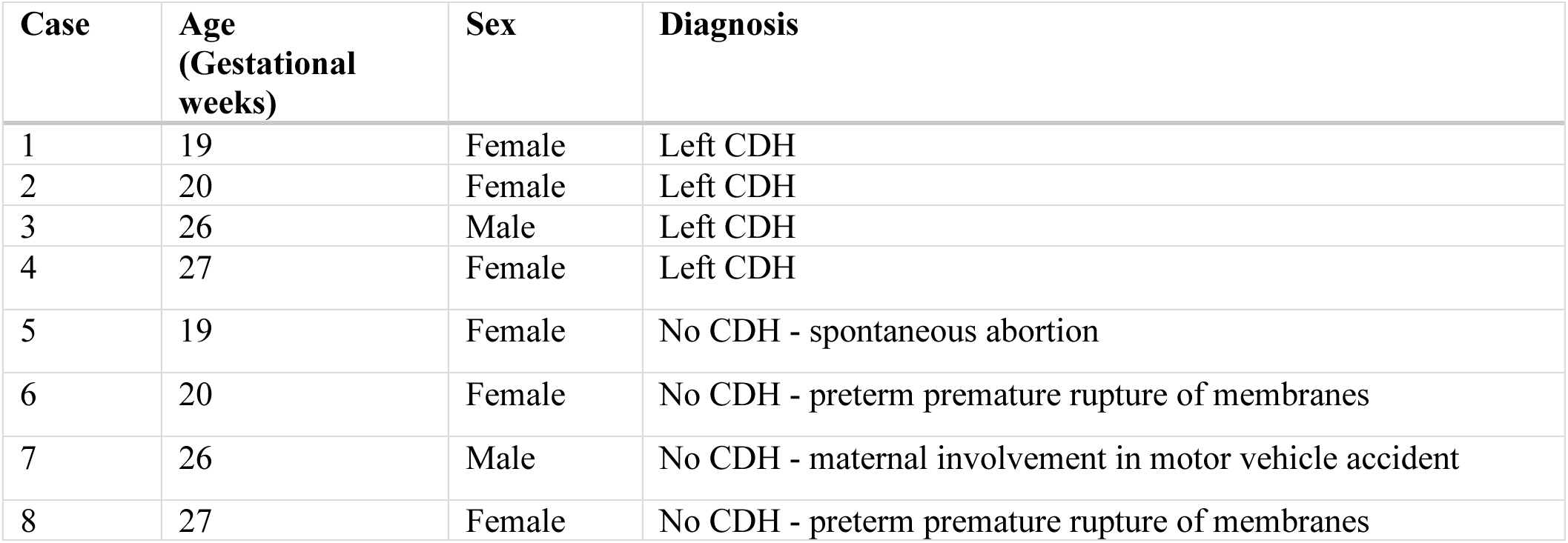
Clinical details of autopsy samples used in this study.

### The severity of cerebral hypoxia in experimental and human CDH is proportional to the degree of pulmonary hypoplasia

To explain the mechanism behind the observed decreased perfusion in the brain of fetuses with CDH, we hypothesized that the compression exerted by the intra-abdominal organs herniated into the chest affected lung and heart morphology. We first confirmed that compression caused by CDH was lungs had lower radial alveolar count (RAC), higher mean liner intercept (MLI), and reduced surfactant production compared to healthy controls and nitrofen-exposed fetuses without CDH (Figure 5A). These findings indicated that the presence of CDH was associated with a higher degree of pulmonary hypoplasia, as established by several research groups in the past (19, 20). We also found a negative correlation between the degree of brain hypoxia, measured as Hif1α expression, and the degree of pulmonary hypoplasia, measured as RAC (Figure 5B) and surfactant protein expression (Figure 5C). Next, we assessed the morphology of the heart in all three conditions. As previously reported (21), we confirmed that nitrofen-exposed fetuses with CDH had left ventricular hypoplasia compared to healthy controls and nitrofen-exposed fetuses without CDH (Figure 5C). These results indicate that CDH is associated with severe lung and left ventricular hypoplasia.

**Figure 5.**
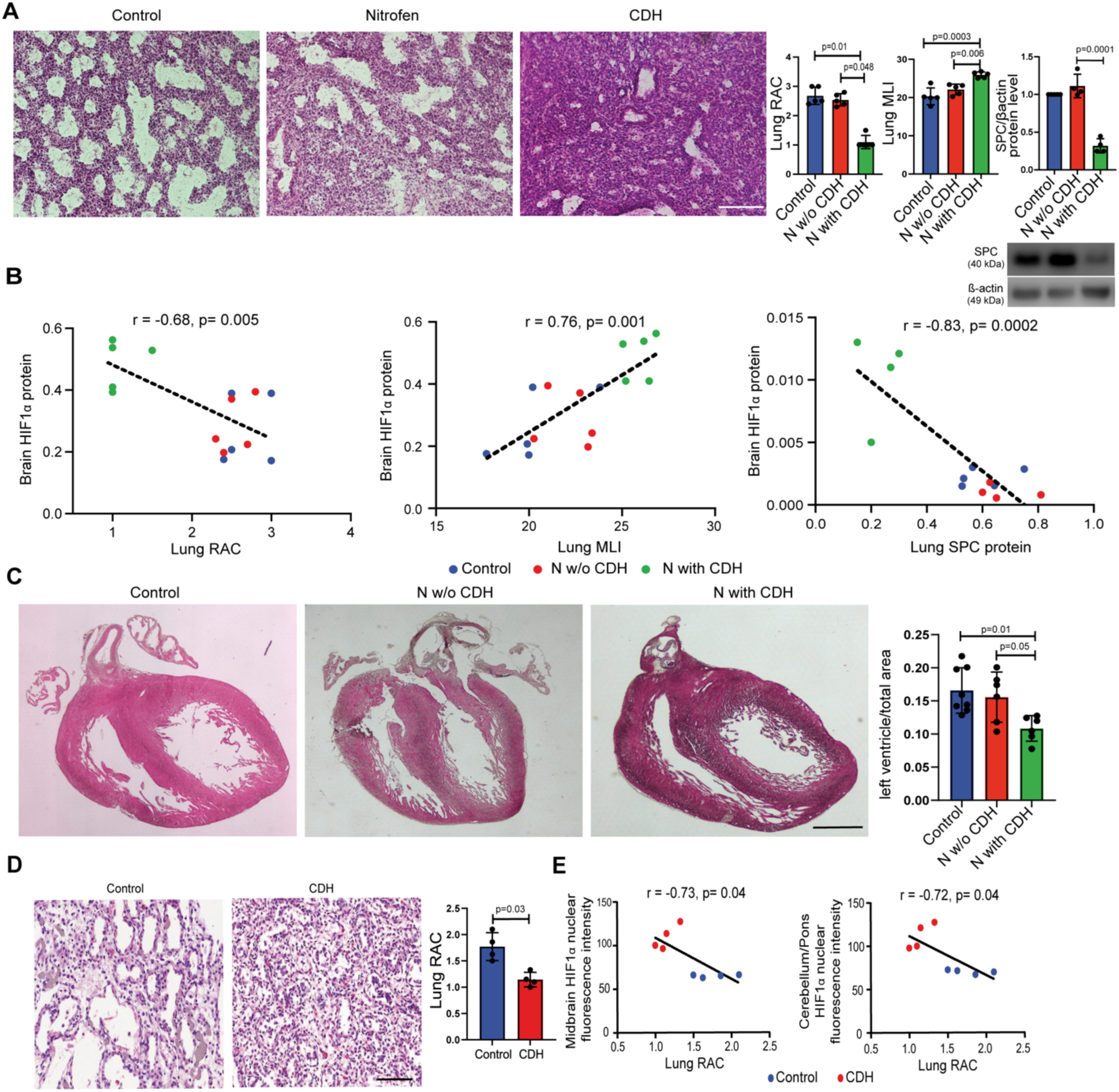
The severity of cerebral hypoxia in experimental and human CDH is proportional to the degree of pulmonary hypoplasia. **A** Representative images of rat fetal lungs stained with hematoxylin and eosin, quantified by radial arterial count and mean linear intercept. Control (n=5), N w/o CDH (n=5), and N with CDH (n=5). Western blot of surfactant protein expression relative to b-actin loading control. Control (n=5), N w/o CDH (n=4), and N with CDH (n=5). **B** Correlation analysis of HIF1α protein expression in the brain and markers of lung maturity in the lung. Pearson correlation statistic is shown. **C** Representative coronal images of the rat fetal heart stained with hematoxylin and eosin, quantified by the ratio of left ventricle to total area. Control (n=8), N w/o CDH (n=6), and N with CDH (n=6). Data are presented as mean ± SD. **D** Representative image of human fetal lungs from CDH and controls, quantified by radial alveolar count (RAC) Scale bar = 50 μm. **E** Correlation analysis between lung RAC and HIF1α fluorescence intensity in the midbrain (right) and cerebellum (middle). Data are presented as mean ± SD.

We also confirmed that the four human fetuses with CDH had pulmonary hypoplasia, as shown by the lower number of airspaces compared to controls (Figure 5D). As observed in the rat model, the severity of pulmonary hypoplasia in human fetuses positively correlated with HIF1α expression in the midbrain and cerebellum (Figure 5E).

## Discussion

This is the first study to show that the brain of experimental and human fetuses with CDH is hypoxic and undergoes cellular and molecular changes that may explain long-term NDI in postnatal survivors. In our study, evidence of a hypoxic state of the brain in CDH fetuses derives from three approaches: imaging, molecular biology, and histology. Using ultrafast power Doppler in fetal rats, we recapitulated the impaired brain perfusion reported in human fetuses with CDH (6–8). This is in line with a recent study employing a rabbit model of CDH that also reported a reduction in MCA flow using conventional Doppler ultrasound (22). In our study, we used ultrafast power Doppler, as this imaging modality allows for a more sensitive detection of blood volume compared to other imaging techniques and has been used in neonates to create high-definition maps of brain vascularization (23, 24). Given its high level of accuracy, one could foresee the use of ultrafast power Doppler extending to clinical application in human CDH fetuses in the future. Further evidence of cerebral hypoxia was shown by the high expression levels of HIF1α, master regulator of hypoxic response, and its hypoxia-inducible target gene, GLUT1. These factors are upregulated in perinatal conditions characterized by either an acute ischemic event, such as in hypoxic ischemic encephalopathy, or by chronic hypoxemia, as in fetal growth restriction (25–27). Lastly, the presence of compensatory angiogenesis in the fetal brain further confirms the hypoxic brain injury observed. Compensatory angiogenesis is a known phenomenon initiated by hypoxia in response to ischemia (28–31) and depends on the activation of factors that we found upregulated in our experiments, such as VEGF, PDGFR, Ang2, and Tie2 (32, 33). New blood vessel formation is also partially regulated by matrix metalloproteinases (MMP) that promote angiogenesis by degrading the extracellular matrix, and by IL-8, a potent pro-angiogenic chemokine that stimulates MMP production (34).

The changes in brain perfusion observed in our rat fetal model of CDH were associated with high levels of apoptosis, a known consequence of brain ischemic insult to both neuronal and glial cells (35). In our study, high levels of apoptosis were found in the stem cell niches of the brain, that is the areas where progenitor cells reside and differentiate into neurons and oligodendrocytes. Neurons and their precursors are highly sensitive to changes in oxygenation and their density is affected in ischemic conditions (36–38). Similarly, mature oligodendrocyte formation from premyelinating OPCs is a critical process for myelination to occur and is known to be affected by prenatal hypoxia (39, 40). Our findings are similar to those reported in the study by Van der Veeken et al, where the brain of fetal rabbits with CDH had similar reduction in neuron density in the hippocampus and decreased OPCs (22). Analogously, experimental studies on a prenatal hypoxemia model in fetal sheep reported changes in brain tissue homeostasis, compensatory angiogenesis, and brain cell population changes, with reduction in neuronal density and myelination integrity (41–44). Overall, the sequalae of reduced brain perfusion in our CDH model are similar to those observed in fetal growth restriction, a condition caused by a chronic reduction in oxygen supply secondary to placental insufficiency (45). In this condition, the brain undergoes progressive neuronal loss, reduction in myelin content, delayed oligodendrocyte maturation, altered vascularization, and an increase in neuroinflammation mediated by activated microglia (45–47). Some studies have speculated that these features are responsible for postnatal NDI in infants born from pregnancies affected by placental insufficiency (36, 48). Similarly, we speculate that postnatal NDI in some CDH survivors may be the result of impaired brain perfusion *in utero*.

To corroborate our experimental findings, we analyzed brain autopsy specimens from human fetuses with CDH who died or were terminated *in utero*. In these specimens, we found that the cerebellum had signs of hypoxia, with upregulation of HIF1α, and associated neuronal and oligodendrocyte loss. These findings are in line with two recent studies that reported alteration in cerebellar morphology in fetuses with CDH (7, 10). Radakrishnan et al reported that magnetic resonance imaging in 63 fetuses with CDH revealed changes in the cerebellum, with specific alterations in vermian dimensions (10). Interestingly, the severity of these changes correlated with the severity of pulmonary hypoplasia. Similarly, Van der Veeken et al retrospectively reviewed the brain ultrasound scans of a large cohort of fetuses with CDH and reported a smaller than average cerebellum in a normal-sized head (7). The authors speculated that these changes in cerebellar dimensions may be attributed to impaired perfusion, diminished venous return, or an underlying genetic cause. Our analysis conducted on brain sections showed upregulation of HIF1α in the cerebellum, thus supporting the impaired perfusion hypothesis.

Our model provides some evidence that the mechanism behind the brain changes herein reported are directly due to the mechanical compression exerted by herniated organs on the mediastinum (Figure 6). This affects not only the lungs, but also the left heart which we found to have changes suggestive of ventricular hypoplasia, as previously reported in human and experimental CDH (49, 50). We speculate that compression of the mediastinal structures results in decreased MCA flow and reduced brain perfusion, with a picture of impaired cerebral oxygenation similar to that seen in fetuses with congenital heart disease (51, 52). This results in increased NPC and OPC apoptosis and decrease in neuronal and oligodendrocyte density. We posit that these series of events could be considered as the first hit to the brain of CDH fetuses that could be followed by a second hit postnatally by invasive maneuvers such as prolonged mechanical ventilation or extracorporeal life support. These changes could potentially explain the predisposition to neurodevelopmental delay that some CDH survivors display later in life.

**Figure 6.**
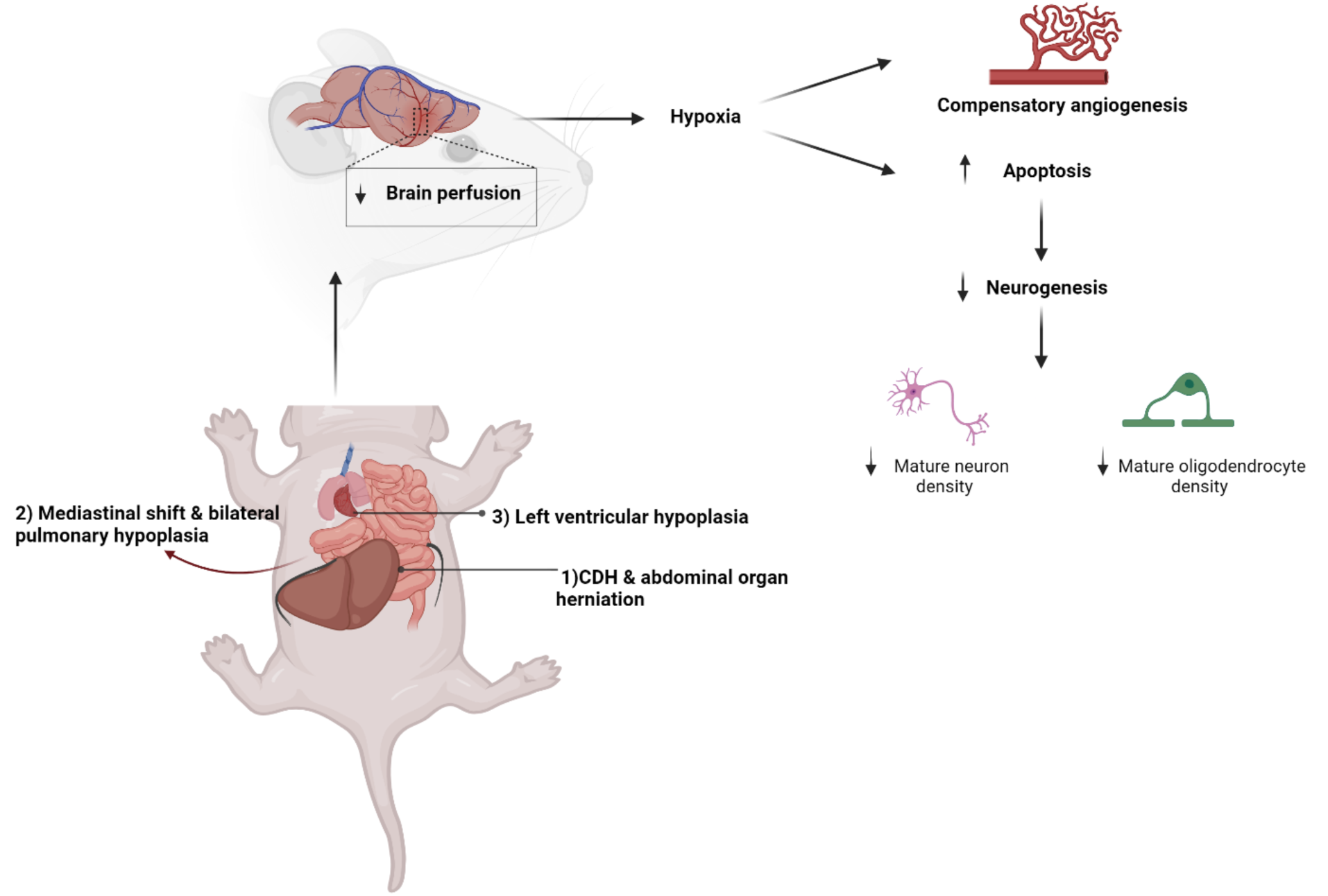
Proposed mechanism of brain injury in CDH. Intrabdominal organ herniation into the mediastinum causes a shift that exerts mechanical compression on the heart and lung causing decreasing cardiac output and pulmonary hypoplasia. This leads to a reduction in brain perfusion, increasing hypoxia signaling and compensatory angiogenesis. The changes in oxygen tension leads to increase apoptosis of neural progenitor cells in the stem cell niches which affects the density of mature neuronal and oligodendrocyte cell types in different regions of the brain.

By characterizing the pathophysiology of brain changes in fetuses with CDH, this study has identified potential targets for treatment. These include the relief of mechanical compression, addressing the hypoxic insult, or improving neurogenesis to rescue brain cell population loss.

Although the brain changes herein described are likely the result of mechanical compression on mediastinal structures, antenatally reducing the herniated organs into the abdomen and closing the diaphragm is not a feasible strategy. In fact, CDH surgical repair *in utero* was attempted in the 1990s but proved to be associated with high rate of fetal demise and was eventually abandoned (53). Fetal endoluminal tracheal occlusion (FETO) is an antenatal intervention that has been shown to increase survival in severe cases of pulmonary hypoplasia secondary to CDH (1, 54). While FETO improves growth and maturation of CDH hypoplastic lungs, this approach does not improve cerebral perfusion (7), likely because the compression exerted by the herniated organs on the mediastinal structures is still present. An alternative to addressing mechanical compression is to reverse brain hypoperfusion. This could be achieved with the use of maternal hyperoxygenation, a strategy that has been investigated to improve fetal developmental outcomes of the lungs, heart, and brain in experimental and human cases of fetal growth restriction and congenital heart disease (55–57). In a study, administration of maternal hyperoxygenation during the third trimester to mothers carrying fetuses with congenital heart disease resulted in increased aortic flow and cerebral oxygen delivery, leading to an improvement in fetal brain growth (58). In experimental models, maternal hyperoxygenation during mid-neurogenesis was shown to increase neural precursor cell density and accelerate cortical development in normal mouse fetuses (58, 59). As maternal hyperoxygenation has proven to be safe for mother and fetus, this approach should be tested in CDH. Finally, a regenerative medicine approach could also prove effective to rescue the loss of brain cell populations. Several preclinical and clinical studies have reported that administration of stem cells or their derivatives have regenerative and neuroprotective effects in fetuses with fetal growth restriction (60, 61). To date, this approach has not been tested in CDH fetuses, likely due to the lack of understanding of the pathophysiology of brain injury, which is now unveiled by the current study.

We acknowledge that our study has some limitations. First, it was not possible to correlate our findings in fetuses with CDH with postnatal neuro-behavioral tests. This was due to the fact that rodent models of CDH are non-surviving, as soon after birth they would require critical care maneuvers and surgery that are not technically feasible. Similarly, we were not able to correlate the human findings with postnatal tests, as our human samples were obtained from autopsy studies. Second, these human samples likely represent the most severe cases of CDH and pulmonary hypoplasia and do not take into account the wide variability that is observed in this disease. Although we do not expect that all fetuses with CDH would have such severe brain changes, our study explored the brain changes in human fetuses that were likely to develop NDI postnatally. Lastly, our study was not able to capture whether the hypoxic brain changes were transient or chronic in CDH fetuses. Longitudinal studies that track brain changes over time in the experimental model are needed to identify the fate of cell populations and the innate responses of the brain.

In conclusion, our study demonstrates that human and rat fetuses with CDH have impaired brain perfusion that leads to an alteration of cellular homeostasis with an increase in apoptosis in the stem cell niches of the brain. This results in a reduction in neural progenitor cells leading to a decrease in mature neurons and oligodendrocytes and associated microglia activation. As CDH fetuses with more severe pulmonary hypoplasia have worse hypoxic brain injury in our study, we speculate that these fetuses are the ones likely predisposed to having long-term NDI. Overall, we propose a two-hit hypothesis that explains NDI in CDH with a first hit characterized by the antenatal loss of neurons and impaired myelination, and a second hit that is caused by postnatal stress factors.

## Methods

### Rat model of CDH

Following ethical approval (Animal Use Protocol #49892), pregnant Sprague Dawley rats were gavage fed nitrofen (100mg) (Sigma Aldrich, USA) to induce fetal pulmonary hypoplasia and CDH, or vehicle alone at embryonic day 9.5 (E9.5) as previously described (12, 13). At E21.5, fetuses were harvested, assessed for presence of CDH, and the brain, lungs, and heart were frozen for RNA/protein analysis or fixed for histological assessment. The following groups of fetuses were compared:

1) control with no pulmonary hypoplasia or CDH;
2) nitrofen exposure with pulmonary hypoplasia without CDH (no compression group, “N w/o CDH”);
3) nitrofen exposure with pulmonary hypoplasia and CDH (compression group, “N with CDH”).

### Autopsy samples

Following ethical approval by the Research Ethics Board at our institution (IRB#1000074888), brain and lung sections from autopsy samples were obtained from 4 CDH fetuses and 4 age- and sex-matched controls (Table 1).

### Ultrafast ultrasound on rat fetal brains

We used an 18Mhz linear array probe (L22-14vX, Verasonics®) attached to a custom fitted portable Verasonics® ultrasound system (Vantage 256, Verasonics Inc.) to obtain brain scans from rat fetuses (**23**). With the rat fetus still attached to maternal-fetal circulation, the ultrasound probe was applied onto the uterus directly on the fetal vertex and held in place during the short time of acquisition. One mid-sagittal acquisition was performed using systematic anatomical landmarks (brain stem and orbits to center along the midline). Singular value decomposition (SVD) filter was applied to the raw ultrafast ultrasound images allowing to distinguish between blood, tissue and noise signal intensities and obtain a power Doppler image (23, 62). A top-hat image processing was applied to the raw Power Doppler images for background suppression. Three anatomical regions of interest (ROI) were selected to approximate the whole brain perfusion. Average signal intensities were extracted to obtain values for whole brain (sum of the 3 regions) and individual regional cerebral blood volume (CBV). Average CBV was calculated for each experimental group. Average CBV of the control group was normalized to 100% and values of nitrofen and CDH groups were individually recorded as percentages compared to the controls.

### Gene expression (RT-qPCR)

RNA was isolated from rat brain tissue using Trizol reagent (TRIzol™ Reagent, Invitrogen™) following the manufacturer’s recommended protocol. Purified RNA was quantified using a NanoDrop™ spectrophotometer (Thermofisher Scientific), and 1 μg of RNA was used for cDNA synthesis (qScript cDNA Supermix, Quantabio). RT-qPCR experiments were conducted with SYBR™ Green Master Mix (Wisent Inc.) for 40 cycles (denaturation: 95 °C, annealing: 58 °C, extension: 72 °C) using the following primer sequences (Supplementary Table 1). Melt curve plots were generated to determine the target specificity of the primers. ΔΔCT method was used to determine normalized relative gene expression of both cytokines.

### Protein expression

Protein lysates were obtained from the rat cerebral cortex, hippocampus, and cerebellum using Tissue Extraction Buffer (Invitrogen™) supplemented with proteinase inhibitors. The total protein yield was determined by Pierce Bradford assay (Pierce™ BCA Protein Assay Kit, ThermoFisher Scientific). Proteins were then transferred to membranes and immunoblotting was performed using the antibodies described in Supplementary Table 2.

### Cerebral vascular density

Vascular density of rat fetal brains was measured by fluorescein isothiocyanate (FITC)-Dextran labelling as previously described (63). Briefly, at E21.5 rat fetuses were placed under isoflurane anesthesia and intracardiac perfusion was performed with PBS to wash out blood components followed by 1mg/ml 10,000 Da FITC (Sigma Aldrich) in PBS (pH 7.0) perfusion for 2 minutes. Rat fetuses were then sacrificed via cervical decapitation, brains were harvested and fixed in 4% paraformaldehyde, and then placed in PBS until imaging. A two-photon confocal laser microscope (Leica) was used to determine vessel density of the whole brain by measuring fluorescence intensity. The microscope was then directly positioned onto the brain dorsolaterally and images were obtained at two fields in the forebrain and hind brain (both left and right side of the brain). Z-stack images were acquired every 5 µm at 20x magnification at a maximum depth of 50 µm. 3D rendering of the images was conducted and total vessel coverage area per field was measured using Volocity software (Quorum Technologies, USA).

### Rat brain morphometric assessment

The head circumference of rat fetuses was measured using a measuring tape and the whole brain was bluntly dissected, harvested, and weighed (Ohaus CS Scale). The fetal rat groups were compared for head circumference, brain weight, and brain to body-to-body weight ratio.

### Apoptosis assessment in stem cell niches

#### TUNEL assay

The brain of rat fetuses from each experimental group were formalin-fixed, paraffin-embedded, and sectioned at 5-μm thickness sagittal orientation in the brain. Following deparaffinization, we stained apoptotic cells using the Click-iT TUNEL kit (ThermoFisher Inc.) according to manufacturer recommended protocols in stem cell niches and in human brain sections. Briefly, deparaffinized sections were post-fixed and de-permeabilized using Proteinase K. Terminal deoxynucleotidyl transferase (TdT) enzyme and 5-Ethynyl-dUTP (EdUTP) nucleotide solution was added to the sections to incorporate EdUTP into double stranded DNA breaks. Following this, a fluorescent azide was added onto sections and EdUTP was fluorescently detected with Click-iT chemistry. After detection, further immunofluorescence co-staining experiments were conducted to mark Dcx+ NPCs, NG2+ OPCs, and nuclei (DAPI). Z-stack images were acquired in the SGZ and SVZ regions using confocal microscopy (Leica SP8).

#### Quantification of TUNEL

3D images were processed on IMARIS 10.1.1 software (Oxford Instruments). We quantified the volume of TUNEL fluorescence intensity specifically on Dcx+ NPCs and NG2+ OPCs along the SGZ and SVZ. Values were averaged between three different sections of the same sample (technical replicates).

### Immunofluorescence experiments

Immunofluorescence assays on rat brain sections were performed by de-paraffinization and antigen retrieval using 10-mM sodium citrate buffer, pH 6. Samples were then incubated with blocking buffer (3% Bovine Serum Albumin (BSA) (Sigma Aldrich, USA) in 1X Tris-buffered saline (TBST) (Wisent Inc.). Primary antibodies were diluted in 1% BSA in TBST, added to the sections, and incubated for 18 hours (Supplementary Table 2). Slides were then washed in TBST, incubated with appropriate secondary antibodies (Invitrogen), DAPI stained (Vector Laboratories Inc), and mounted.

#### Fetal rat brain

Sections of rat fetal brain from all groups were used to determine the cellular density of the following populations using immunofluorescence: mature neurons (NeuN), oligodendrocytes (Olig2, Ki67), and activated microglia (TMEM119). Individual fields within the cerebral cortex, hippocampus, and cerebellum were acquired at 20x magnification in Pannoramic Viewer (3D Histech) imaging software. Quantitative analysis was conducted using a semi-automatic image analysis method in ImageJ (64).

#### Fetal human brain

We obtained sections of human fetal midbrain, cerebellum, and hippocampus from autopsy samples (Table 1). We determined the cellular density of the following populations using immunofluorescence: mature neurons (Hu/CD), oligodendrocytes (progenitors: NG2 and mature: Olig2 and MBP), activated microglia (Iba1 density and cell morphology) and astrocytes (GFAP) (Supplementary Table 2). Confocal laser microscopy (Leica SP8) was used to image at least 5 individual fields at 40x magnification per sample. Density of positive cells was quantified using the same method described above for the rat fetal brain.

### Lipid peroxidation assay

Protein lysates were used to conduct lipid peroxidation assay using colorimetry (Lipid Peroxidation, MDA Assay kit (abcam) following manufacturer’s recommended protocol. Absorbance was measured using a microplate reader at 532 nm OD and the concentration of MDA in each sample was determined.

### Lung and heart histology

#### Rat lung

The left lungs of fetuses were harvested from each experimental group and flash frozen for cryo-sectioning. Lungs were embedded in Optimal Cutting Compound, sectioned at 5-μm thickness, stained using hematoxylin and eosin (H&E), dehydrated, and then mounted in xylene based mounting media. Brightfield microscopy (Olympus) was used in all groups (n=6, 3 sections per sample per marker) to analyze lung morphometry by RAC and MLI as previously described (65).

#### Rat heart

Coronal sections of hearts were stained and imaged using brightfield microscopy (Olympus) at 20x magnification. To assess the degree of left ventricular hypoplasia, the ratio of the area of the left ventricle and the total area of the heart was measured on Image J.

#### Human fetal lung

Sections from the left lung of human fetal autopsy samples were H&E stained and assessed for severity of pulmonary hypoplasia by RAC and MLI, in accordance with the guidelines of the American Thoracic Society (65).

### Statistics

Groups were compared using the appropriate parametric and non-parametric testing including Student t-test, one-way ANOVA (Tukey post-test), or Kruskal-Wallis (post-hoc Dunn’s non-parametric comparison) test according to the Gaussian distribution assessed by Shapiro Wilks normality test. Correlation analysis was conducted by Pearson correlation and simple linear regression. P value was considered significant when p<0.05. For ANOVA comparisons, appropriate F and H values are presented in the figure legends. Furthermore, for unpaired and paired Student t-tests, appropriate Man-Whitney U value and a T or W (Wilcoxon) value respectively are reported in the figure legends. All statistical analyses were produced using GraphPad Prism® software version 10.0.

### Study approval

The animal studies were approved by our animal care committee (AUP #49892). Human autopsy samples were obtained following approval form institutional review board (IRB#1000074888)

## Data availability

Values for all data points in graphs are reported in Supporting Data Values in file.

## Author contributions

GB, LA, AZ designed the research studies. GB, JA, KK conducted experiments. GB, LA, NF, JB, analyzed the data. GB, LA, AZ wrote the manuscript, and LM, OV critically appraised the manuscript.

## Acknowledgements

The authors would like to thank Rachel Bervocitch, Deivid Rodrigues, Jeanette Reyes, David Chiasson, Gino Somers, Lab Animal Service, and Imaging Facility at the Hospital for Sick Children for their support of this project. This study was supported by the Canadian Institutes of Health Research (CIHR) Project Grant (175300), Perioperative Services Facilitator Grant at the Hospital for Sick Children, Toronto, and the SickKids Congenital Diaphragmatic Hernia Fund (R00DH00000), and Natural Sciences and Engineering Research Council of Canada (NSERC), Canada (RGPIN-2021-03539).

---

**Supplementary Figure 1.**
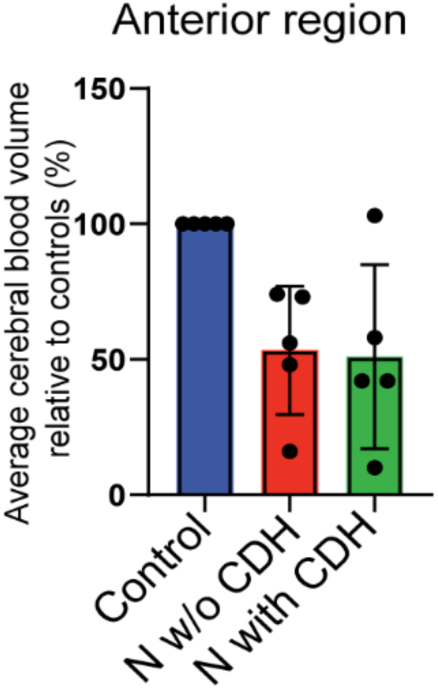
Cerebral perfusion quantification of the anterior brain region in CDH fetal rats. No differences in cerebral perfusion were observed among three experimental groups (. Data is presented as mean ± SD.

**Supplementary Figure 2.**
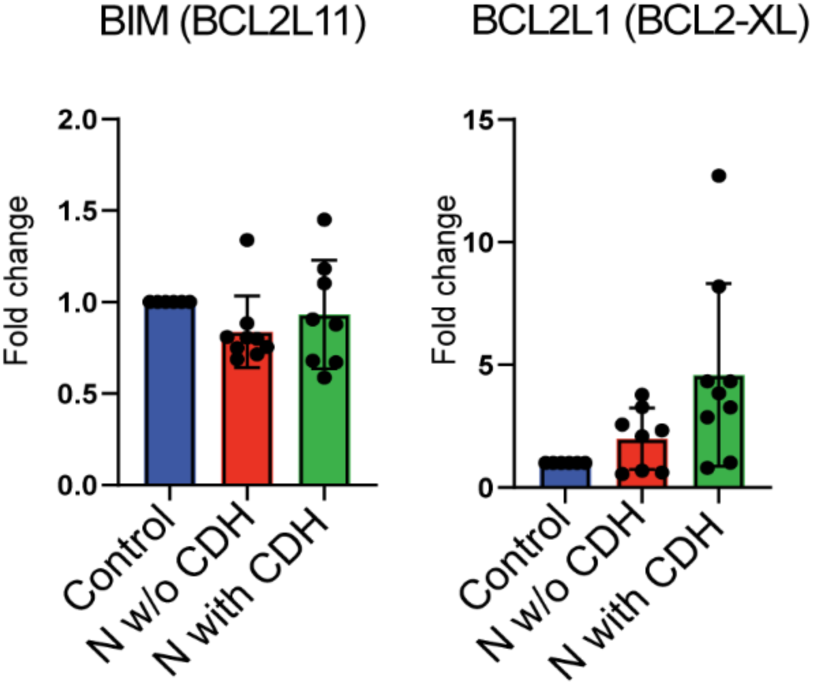
Gene expression of anti-apoptotic marker. RT-qPCR experiments show no differences in gene expression of anti-apoptotic *BCL2L11* or *BCL2-XL* in the brain of pups among the three experimental groups. Control (n=6), N w/o CDH (n=8), N with CDH (n=7). Data are presented as mean ± SD.

**Supplementary Figure 3.**
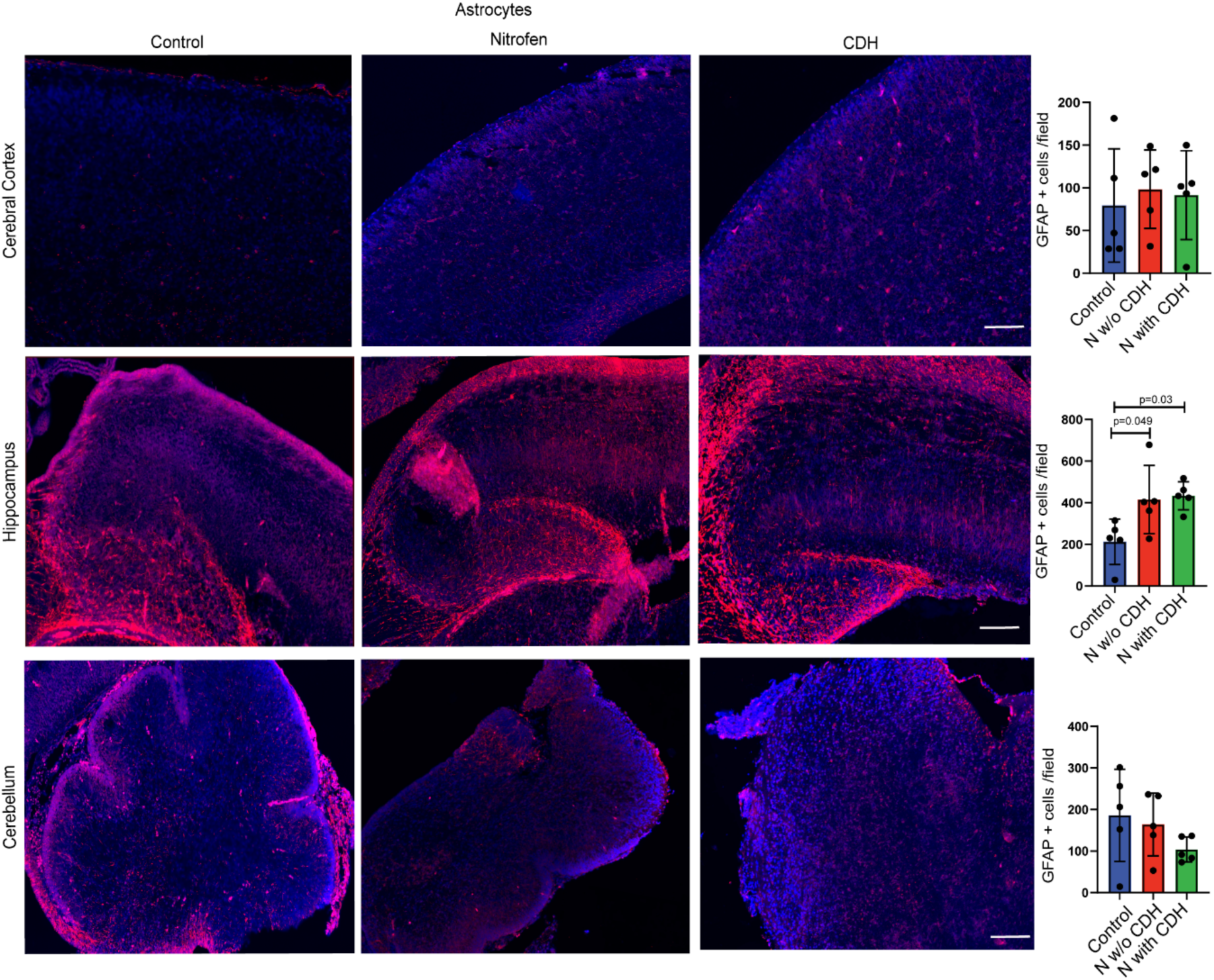
Astrocyte activation in the CDH rat fetal brain. Representative immunofluorescence images of astrocyte marker glial fibrillary acidic protein (GFAP, red) and nuclear marker (DAPI, blue) in the cerebral cortex, hippocampus, and cerebellum. Nitrofen-exposure alone caused an increase in astrocyte activation in the hippocampus. Control (n=5), N w/o CDH (n=5), N with CDH (n=5). Data are presented as mean ± SD. Scale bar = 50 μm.

**Supplementary Figure 4.**
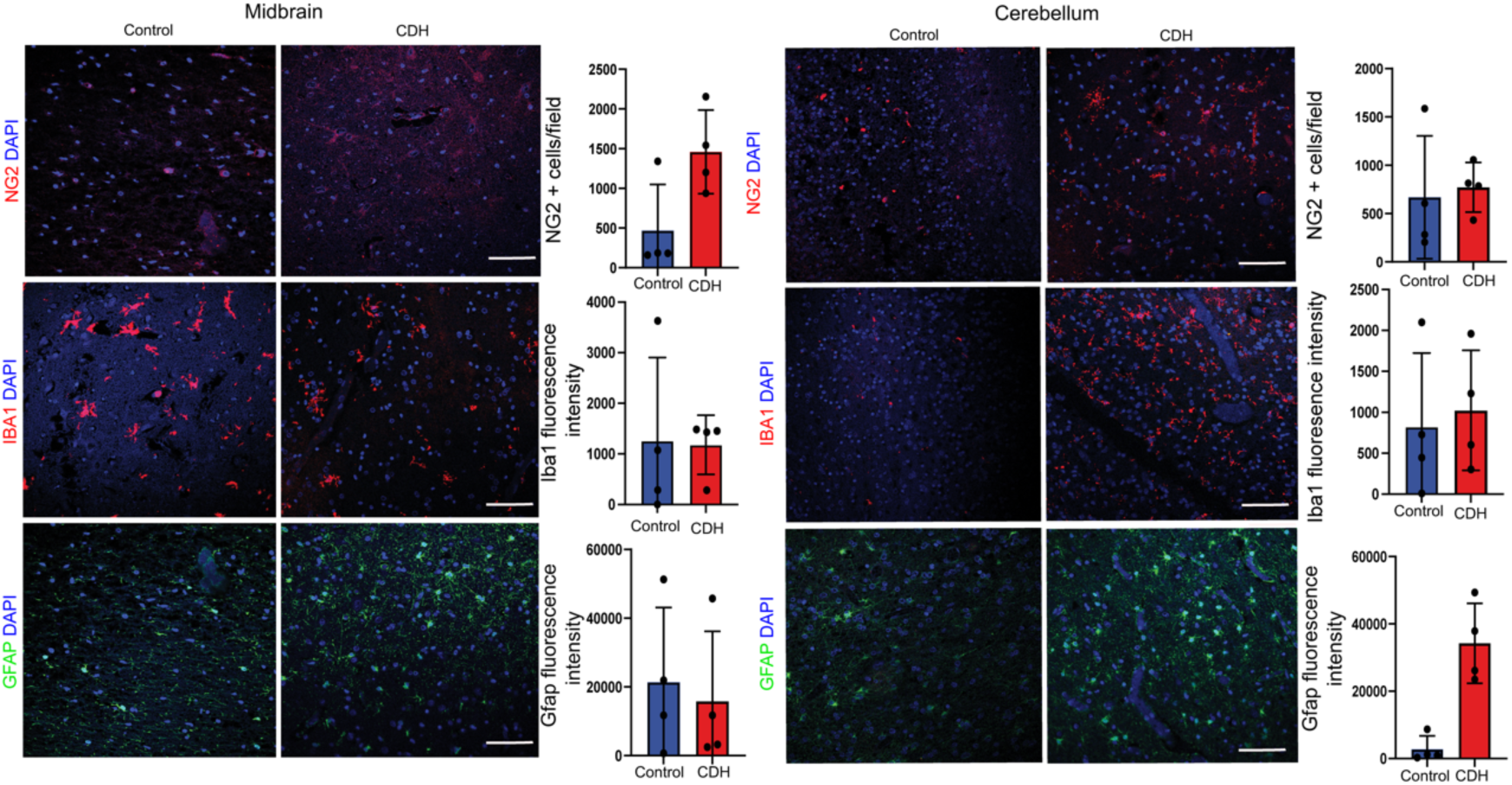
Oligodendrocyte progenitor cells, microglia, and astrocytes in the midbrain and cerebellum/pons of human fetuses. Representative immunofluorescence images of oligodendrocyte progenitor cells (NG2+) activated microglia (IBA1+), and astrocytes (GFAP+), quantified by number of cells per field (NG2) or fluorescence intensity (Iba1, GFAP).

**Supplementary Figure 5.**
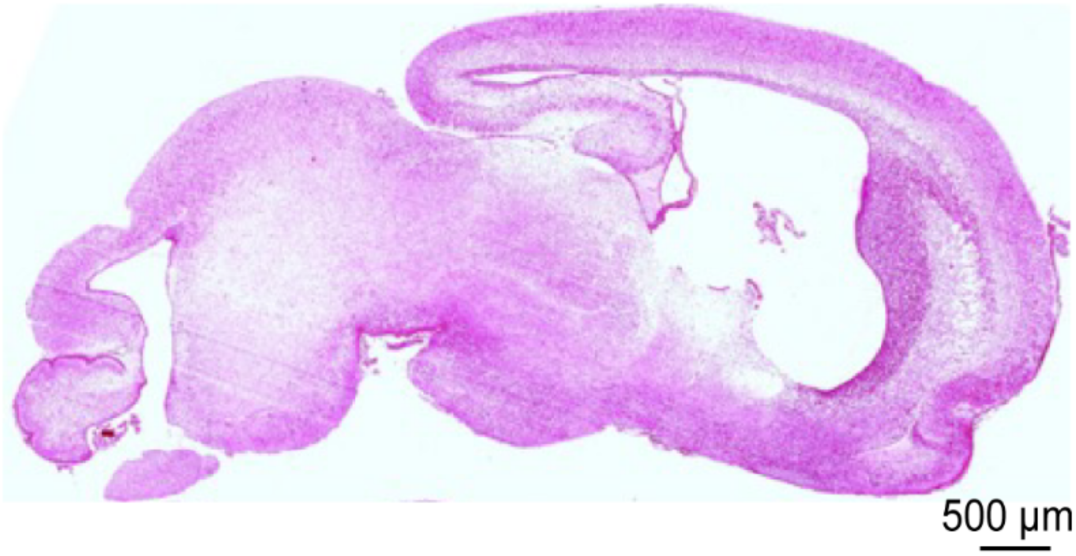
Representative image of Hematoxylin and Eosin stained sagittal/medial brain section of E21.5 rat pup at the specific histological level used for the immunofluorescence experiments.

**Supplementary Table 1.**
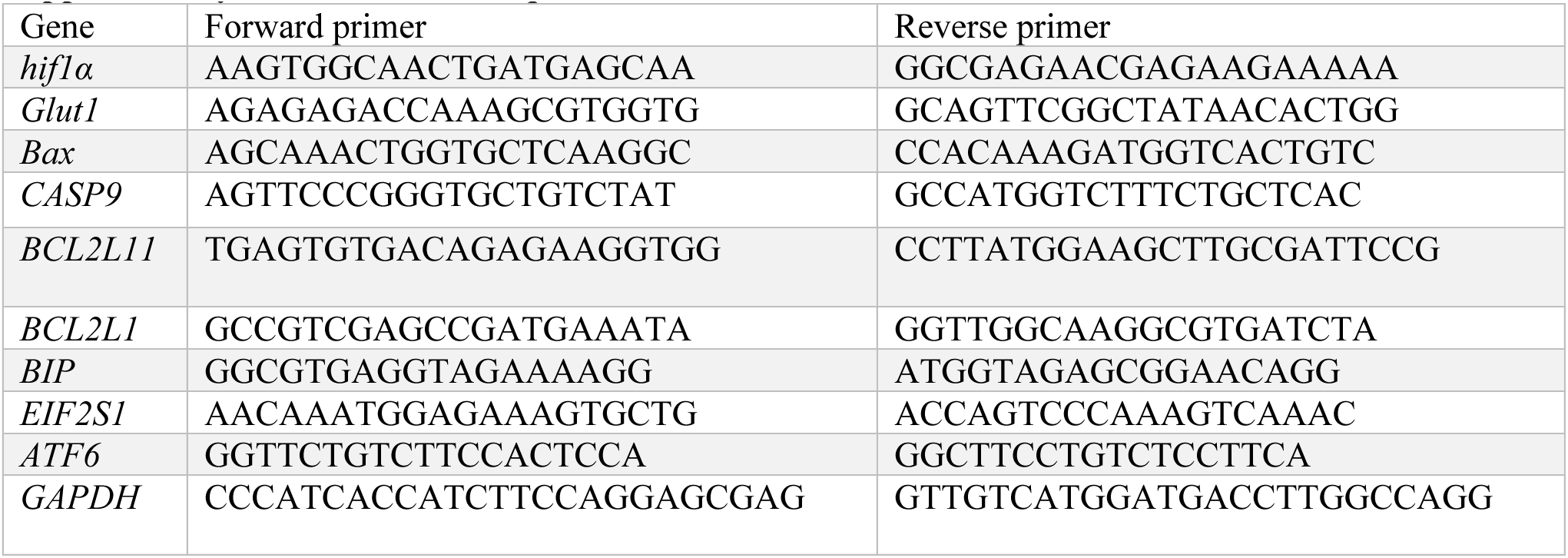
Primer sequences.

**Supplementary Table 2.** Primary & Secondary antibodies.

| Primary antibody | Marker | Species | Assay & Dilution | Company | Catalog number |
| --- | --- | --- | --- | --- | --- |
| HIF1α | Hypoxia | mouse | WB: 1:1000<br>IF: 1:100 | Novus | NB100-105 |
| Glut1 | Hypoxia | rabbit | WB: 1:1000 | Santa Cruz | sc-7903 |
| Histone 3 | Loading control | rabbit | WB:1000 | Cell signaling | 9715S |
| Bax | Pro-apoptosis | rabbit | WB: 1:1000 | Cell Signaling | 2772 |
| CC9 | Pro-apoptosis | rabbit | WB: 1:1000 | Cell Signaling | 9509P |
| BiP | ER stress | rabbit | WB: 1:1000 | Cell Signaling | 3177T |
| p-eIF2α | ER stress | rabbit | WB: 1:1000 | Cell signaling | 9721S |
| ATF6 | ER stress | mouse | WB: 1:1000 | Novus | NBP1-40256SS |
| NeuN | Mature neurons | mouse | IF: 1:200 | EMD<br>Millipore<br>Sigma | MAB377 |
| TMEM119 | Microglia | rabbit | IF: 1:100 | abcam | ab185337 |
| Ki67 | Proliferation | rat | IF 1:200 | Novus<br>Biologicals | NB500-170 |
| Olig2 | Oligodendrocytes<br>(progenitor and<br>mature) | mouse | IF: 1:50 | R and D<br>systems | AF2418 |
| NG2 | Oligodendrocyte<br>progenitor cell | rabbit | IF: 1:200 | abcam | ab275024 |
| MBP | Myelin basic protein | rabbit | IF: 1:200 | EDM<br>millipore<br>Sigma | M3821 |
| GFAP | Astrocytes | chicken | IF: 1:500 | abcam | ab4674 |

